# Superposition of survival curves as a tool for epistasis analysis of longevity interventions

**DOI:** 10.1101/147173

**Authors:** Stefan Nowak, Johannes Neidhart, Jonas Rzezonka, Ivan G. Szendro, Rahul Marathe, Joachim Krug

## Abstract

A long-standing problem in ageing research is to understand how different factors contributing to longevity should be expected to act in combination under the assumption that they are independent. Standard epistasis analysis compares the extension of mean lifespan achieved by a combination of interventions to the prediction under an additive or multiplicative null model, but neither model is fundamentally justified. Moreover, the target of longevity interventions is not mean life span but the entire survival curve. Here we formulate superposition principles that predict the survival curve resulting from a combination of two interventions based on the survival curves of the individual treatments, and quantify epistasis as the deviation from this prediction. We test the method on a published data set comprising survival curves for all combinations of 4 different longevity interventions in *Caenorhabditis elegans*. We find that epistasis is generally weak even when the standard analysis indicates otherwise.

---

Research on the biology of ageing has revealed a large variety of genetic, metabolic and environmental interventions that increase life span in model organisms [1–5]. Some interventions, such as dietary restriction, are remarkably universal and apply in similar form across widely different species [6, 7]. An important tool that is used to unravel the underlying mechanisms is epistasis analysis, where the effect of a given intervention on lifespan is probed in the presence of another manipulation [7–10]. The term epistasis commonly refers to interactions between genetic mutations [11–15] but will be used here in a broader sense that includes also physiological interventions. The interpretation of epistasis studies is relatively straightforward if the effect of the first intervention either persists unchanged or is completely masked by the second. However, in many cases the mutual influence of different interventions is quantitative rather than qualitative, and correspondingly a quantitative criterion of independence is required in order to infer whether and how the interventions interact.

In the past, most studies have focused on mean or median life span as the primary longevity phenotype, employing a plausible null model [16] where either the absolute life span extensions caused by independent interventions are assumed to add up (*additive model*), or the relative increases are assumed to multiply (*multiplicative model*). No clear preference for either of the two null models can be derived from first principles, and therefore it has been recommended that both the additive and multiplicative scales should be used to test for epistasis in longevity data [9]. More importantly, the restriction to mean life span for the quantification of longevity effects neglects the entire information contained in the shape of the survival curve [17–19]. Many studies have incorporated shape information by fitting experimental survival curves to mathematical models [20–27], but only rarely has this approach been used to analyze epistatic interactions in terms of model parameters such as the rate of mortality acceleration [10]. A framework for epistasis analysis that is based directly on the survival curve does not seem to have been proposed previously.

For the following discussion, a survival curve *S*(*x*) is a monotonically decreasing function that quantifies the fraction of the population that is still alive at time x. Accordingly, *S*(*x*) is restricted to the interval [0,1] with limits *S*(0) = 1 and *S*(*x* → ∞) = 0. Then, the purpose of this paper is to address the following question: Given a baseline survival curve *S*_0_(*x*) and survival curves *S*_1_(*x*) and *S*_2_(*x*) resulting from two different longevity interventions, can one predict the survival curve *S*_12_(*x*) that would result if the two interventions acted in combination and independently? We propose several possible answers to this question that are based on different assumptions about the meaning of independence, and which we collectively refer to as *superposition principles* (*SP’s*).

Adopting the view that epistasis, in the most general sense of the term, expresses “our surprise at the phenotype when mutations are combined, given the constituent mutations’ individual effects” [15], the validity of a SP implies the absence of epistasis on the level of the survival curves. Correspondingly, the deviation of the data from the prediction of the SP’s can be used to quantify the amount of epistasis. The implementation of this idea requires to formalize the effect of a given longevity intervention as a mathematical transformation acting on the set of survival curves. As a simple example, consider the temporal rescaling operation *S*(*x*) → *S*(*bx*), where *b* < 1 if life span is increased [28]. If *S*_1_(*x*) and *S*_2_(*x*) arise from the baseline survival curve *S_0_(x)* by temporal rescaling with factors *b*_1_ and *b*_2_, respectively, then the natural prediction for the survival curve of the combined intervention, under the assumption that the two interventions do not interact, is given by

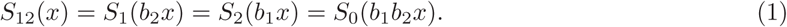

Note that the empirical validity of this relation is far from obvious, even if all four survival curves are indeed related by temporal scaling. In practice, we have found that simple rescaling is generally too restrictive to allow for an accurate description of empirical data. Below we therefore complement the scaling parameters *b*_*i*_ by a second parameter affecting also the shape of the survival curve, and refer to the resulting SP as *generalized scaling SP* or *GS-SP*.

Whereas the implementation of the GS-SP requires to explicitly estimate the parameters defining the transformations leading from *S*_0_ to *S*_1_ and *S*_2_, the other two SP’s are non-parametric. The first is a generalization of the multiplicative null model, which extends the assumption that the relative increases of mean life span combine multiplicatively to the entire quantile function *Q*(*s*). Here *Q*(*s*) denotes the inverse function of the survival curve *S*(*x*), i.e. *Q*(*s*) is the age at which a fraction *s* of the population is still alive. In particular, the median life span is given by *Q*(1/2), and the *generalized multiplicative SP (GM-SP)* reads

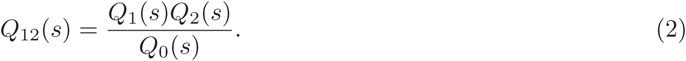

The temporal scaling relation (1) constitutes a special case of (2), and we will see below that the transformations underlying the GM-SP can be viewed as inhomogeneous temporal rescalings where the scale factor depends on the fraction of surviving individuals.

In contrast to the GM-SP, which is motivated primarily by formal considerations, the third SP is based on a clear biological picture and can be formally derived within the reliability theory of ageing [29, 30]. The key assumption taken from this theory is that the survival of an organism requires the maintenance of several vital functional modules, and the organism dies when one of these modules fails. In the language of failure time analysis the failures of different modules are *competing risks* [31], and independence of longevity interventions implies that they affect disjoint sets of functional modules. A straightforward derivation given below then yields the *competing risks SP (CR-SP)*

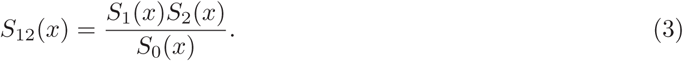

Despite the formal similarity between (2) and (3), their implications are markedly different. First, whereas by construction median life spans combine multiplicatively under the GM-SP (2), and hence standard analysis would detect no epistasis, we will show below that the CR-SP (3) contains a generic mechanism for synergistic epistasis on the level of mean life span. Second, the requirement that the CR-SP yields a valid combined survival curve *S*_12_(*x*) poses rather restrictive conditions on the shapes of the survival curves *S*_0_, *S*_1_ and *S*_2_. By contrast, the GM-SP (2) is more easily satisfied.

After exploring the mathematical properties of the proposed SP’s and discussing their relation to conventional epistasis analysis, we apply them to a published data set containing measured survival curves for all combinations of 4 different longevity interventions in *C. elegans*, viz. two genetic mutations, dietary restriction and cold temperature [10]. As each of the 6 pairs of interventions can occur on 4 different backgrounds, this data set allows for a total of 24 pairwise analyses. For each pair of interventions, we determine parametrized fits to the four survival curves that are constrained to conform to the SP’s and compare them to unconstrained fits. The relative improvement in the accuracy of the fit that is achieved by lifting the constraint can then be interpreted as a measure for the deviation from the specified type of independence. Somewhat surprisingly, we find that most pairs of interventions can be well described by at least one of the SP’s, indicating that the level of epistasis, in the general sense defined above, is low. By focusing on cases where one of the possible fits is significantly better than the others, we identify several characteristic patterns that may provide the basis for a classification of the effect of different longevity interventions on the survival curves. Some general conclusions and open problems for future work that follow from our study are outlined in the Discussion.

## Theory

### Superposition principles

Let *S*_0_, *S*_1_, *S*_2_ and *S*_12_ be a quadruple of survival curves corresponding to two different interventions, i.e., *S*_1_ and *S*_2_ result from *S*_0_ by single interventions and *S*_12_ results from *S*_0_ by combining both interventions. We say that this quadruple fulfills a SP if there are mappings *T*_1_ and *T*_2_ from the set of survival curves onto itself such that

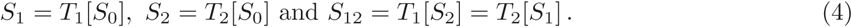

This definition is based on the assumption that longevity interventions can be formally separated from the ageing phenotype on which they act, and that the latter is sufficiently well represented by the survival curve for this approach to be predictive. Although neither of these assumptions is self-evident, the specific examples to be discussed in the following show that the abstract superposition principle (4) unifies several natural conceptualizations of the independence between longevity interventions. It thus provides a useful framework for a generalized, quantitative epistasis analysis on the level of survival curves.

The three specific examples of SP’s described below are not exhaustive, and indeed it appears to be a major mathematical challenge to classify all possible transformations *T*_*i*_ and functions *S*_*i*_ for which (4) holds. However, the logic of our approach only requires to find one SP that is approximately satisfied for a given pair of interventions in order to conclude that epistasis, in the broad sense defined here, is absent or at least weak. Finding the specific SP that minimizes the deviation between the non-epistatic prediction and the data for a particular case is analogous to (but more complex than) the identification of the proper non-linear scale on which to measure a phenotype in order to obtain an unbiased estimate of genotypic epistasis [11, 12, 14, 32].

### Generalized multiplicative SP

The basic assumption underlying this principle is that the age *x = Q*(*s*) at which a certain fraction *s* of individuals is still alive is multiplied by an s-dependent factor *f*_*i*_(*s*) in the presence of an intervention *i*. In particular, the median lifespan *m*_0_ = *Q*_0_(1/2) of the baseline population would be multiplied by *f*_*i*_(1/2). In terms of survival curves, the intervention results in the multiplication of the inverse survival curve with the function *f*_*i*_, i.e., *T*_*i*_[*S*] *=* (S^−1^ *f*_*i*_)^−1^, where *F*^−1^ denotes the inverse of a function *F*. The survival curve corresponding to the double-intervention can then be written as

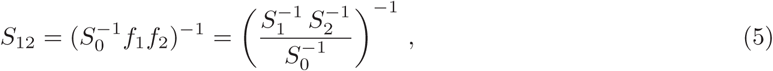

which is equivalent to (2). By construction, validity of the GM-SP ensures that median life spans combine multiplicatively,

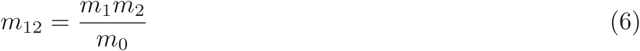

Where *m*_i_=*Q*_i_(1/2).

Equation (2) implies that the GM-SP is fulfilled if *S*_0_, *S*_1_ and *S*_2_ are chosen arbitrarily and *S*_12_ is constructed according to the right-hand side of (5). Note, however, that depending on the choice of the *S*_*i*_ the resulting curve *S*_12_ may not be a valid survival curve. As an illustrative example, consider Weibull survival curves of the form

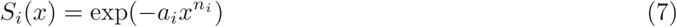

with positive parameters *a*_*i*_ and *n*_*i*_ for *i* = 0,1,2. The inverse function reads *Q*_*i*_(*s*) = (−log(*s*)/*a*_*i*_)^1/*n*_*i*_^ and after some algebra one finds that the combined survival curve is again of Weibull form, *S*_12_ = exp(−a_12_*x*^n_12_^), with 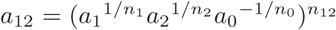 and 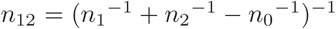. Since *S*_12_ is a valid survival curve only if *n*_12_ > 0, the condition on the parameters is 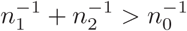.

### Competing risks SP

The reliability theory of ageing [29, 30] uses concepts that were developed in engineering and product design to describe the failure of artificial systems and applies them to living organisms. The basic idea is that the system can be reduced to a series of blocks, where each block consists of parallel redundant elements, and each element has a certain (constant) failure rate. The blocks in series are interpreted as essential functional modules of an organism, such as organs, which consist of redundant elements, such as cells and pathways. Modules cease to function if all their redundant elements have failed, and the death of the organism is caused by the failure of one of the essential modules.

The key feature of reliability theory that is relevant in the present context is that the probability for the organism to survive up to age *x*, that is, the survival curve *S*(*x*), is equal to the product of the probabilities that each of the essential modules is still functional at time *x*. When there are *N* independent modules each characterized by a survival probability *P*_*k*_(*x*), the resulting survival curve has the form

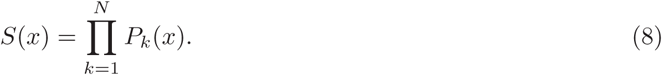

This mathematical structure is known from failure time analysis as a competing risks model [31]. In this setting the failure of each module *k* is a latent cause of death with its own survivor function *P*_*k*_, and the actual time of death or failure is the smallest among the *N* latent failure times. Equation (8) then follows if the risks are independent.

Assuming that a given intervention affects only one of the *N* modules, the corresponding survival probability *P*_*k*_(*x*) is replaced by another function 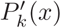, which implies the transformation *T*_*k*_[*S*] = *Sϕ*_*k*_ with 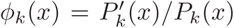. When two interventions affect different modules, the survival curve corresponding to the combined intervention is then indeed given by *S*_12_ = *S*_0_*ϕ*_1_*ϕ*_2_ = *S*_1_*S*_2_/*S*_0_. If each of the focal interventions affects several of the modules the CR-SP remains valid provided the two sets of affected modules are disjoint. It is implicit in the product form of (8) that the modules affected by the two interventions are then not only independent of each other, but also independent of all other determinants of life span that remain unaffected.

The example of Weibull-type survival curves (7) reveals that the validity of the CR-SP (3) places rather restrictive conditions on the parameters. Constructing the double-intervention survival curve yields

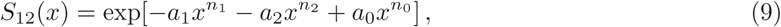

and it is easy to see that a necessary condition for the combined curve to be monotonically decreasing is that min[*n*_1_,*n*_2_] < *n*_0_ < max[*n*_1_,*n*_2_]. Setting *n*_0_ = *n*_1_ = *n*_2_ = *n* the combined survival curve is again of Weibull form, but it is valid only if *a*_0_<*a*_1_ + *a*_2_. In terms of the median life spans this condition reads

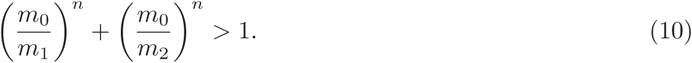

If both interventions are of equal effect, *m*_1_/*m*_0_ ≈ *m*_2_/*m*_0_, this condition can be satisfied only if this effect is rather weak, *m*_1_/*m*_0_ < 2^1/*n*^ where often *n* ≫ 1 [22]. On the other hand, the condition (10) can also be satisfied by interventions of widely different effects, e.g. *m*_1_/*m*_0_ ≈ 1 and *m*_2_/*m*_0_ ≫ 1. When the condition (10) is satisfied, the median life span of the combined intervention is given by

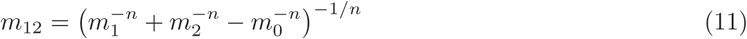

which can be shown to always exceed the multiplicative expectation (6) (see Supplementary Material for the derivation). Thus, at least for the simple case of Weibull survival curves with equal index n, the CR-SP predicts positive (synergistic) epistasis for median life spans, and it is expected to hold preferentially for interventions of strongly unequal effect. We will see below that this pattern is indeed found in empirical data.

Finally, we note that the CR-SP takes a simple form when written in terms of the age-dependent mortalities or hazard rates defined by *h*_*i*_ = −(1/*S*_*i*_)*dS*_*i*_/*dx*. Indeed, using (3) it follows that *h*_12_ =*h*_1_ + *h*_2_ − *h*_0_ or

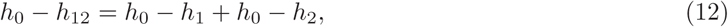

which implies that the reductions of mortality afforded by the two focal interventions add up in the combination.

### Generalized scaling SP

Rather than multiplying a survival curve or its inverse with a function, one can also think of applying a function to a survival curve *S*(*x*) (outer scaling) or to its argument *x* (temporal scaling). This yields a transformation of the general form *T*_*i*_[*S*](*x*) = *g*_*i*_(*S*(*t*_*i*_(*x*))). In order to ensure the validity of the general SP (4), the functions *g*_*i*_ and *t*_*i*_ have to fulfill the conditions *g*_1_(*g*_2_(*x*)) = *g*_2_(*g*_1_(*x*)) and *t*_1_(*t*_2_(*x*)) = *t*_2_(*t*_1_(*x*)), respectively, for all *x*. Furthermore, the functions have to preserve the survival curve properties and hence *g*_*i*_(0) = *t*_*i*_(0) = 0, *g*_*i*_(1) = 1 and *t*_*i*_(x → ∞) = ∞.

A simple choice that satisfies all these conditions is a linear scaling of time [28], *t*_*i*_(*x*) = *b*_*i*_*x*, and a power function applied to the survival curve, *g*_*i*_(*s*) *=s^q_i_^*, with positive constants *b*_*i*_ and *q*_*i*_. Starting from a baseline survival curve *S*_0_, the single-intervention curves are then of the form

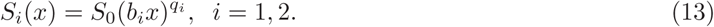

In terms of the hazard rates Eq. (13) takes the form *h*_*i*_(*x*) = *q*_*i*_*b*_*i*_*h*_0_(*b*_*i*_*x*), which shows that the transformation combines an accelerated failure rate model (parametrized by *b*_*i*_) with a proportionate risk model (parametrized by *q*_*i*_) [28,31]. The *generalized scaling SP (GS-SP)* is satisfied if constants *b*_1_, *b*_2_, *q*_1_, *q*_2_ can be found such that the survival curve of the combined intervention is given by

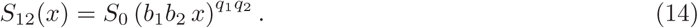

As was mentioned above, in the case of purely temporal scaling (*q*_1_ = *q*_2_ = 1) the transformed curves satisfy the GM-SP (2), but in general the GS-SP does not reduce to any of the other two SP’s. For the special case when *S*_0_ is of Weibull form (7) the transformation (13) amounts to a pure temporal rescaling with scale factor 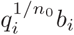, and correspondingly the median life spans combine multiplicatively as in (6), under the GS-SP. However, for survival curves of Gompertz form the GS-SP is consistent with both antagonistic and synergistic epistasis on the level of mean life span (see Supplementary Material for details).

## Data analysis

### Data set

As an illustration of our approach, we analyzed a published data set for *C. elegans* exposed to four different longevity interventions [10]. These included two genetic mutations (*clk-1* and *daf-2*), cold temperature (16° C vs. 25° C at control conditions) and dietary restriction (axenic medium). Survival curves were obtained in triplicate for each of the 2^4^ = 16 possible combinations of interventions. In order to achieve the large cohort sizes required for a meaningful fit of survivorship data to survival functions [21,22], we pooled the replicates for each set of conditions, which yields cohorts of more than 300 individuals. Since each of the 6 pairs of interventions can be applied to 4 different baseline conditions including 0, 1 or 2 other interventions, the data allow for 24 different pairwise comparisons. Each comparison makes use of a quadruple of survival curves comprising the baseline condition, each of the focal interventions applied individually, and the combination of the two focal interventions.

For a better overview of the relation between survival curves, we assign a binary string to each of them. A position of the string corresponds to a certain intervention, with a 0/1 at this position determining whether the corresponding intervention takes place. The assignement of interventions is as follows: The first position indicates reduced temperature, the second the *daf-2* mutation, the third the *clk-1* mutation and the fourth position corresponds to dietary restriction. For example, the string 1001 labels the survival curve at 16°C with dietary restriction but in the absence of genetic mutations. In this notation, a quadruple of survival curves is represented by two strings that differ at two positions and the two intermediate strings that differ in one position from either of the two aforementioned strings. A valid quadruple would be, for instance, 1001 (baseline), 1101, 1011, and 1111. For the sake of brevity we will write 1001–1111 for this quadruple of survival curves. The full list of combinations of interventions is given in Tab. 1.

**Table 1.**
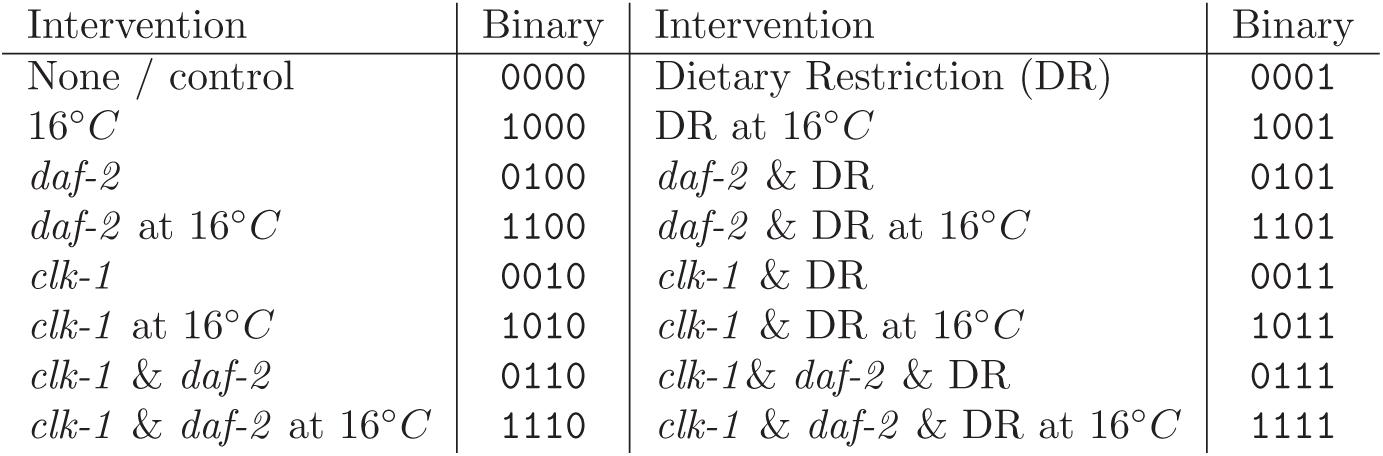
Binary representation used to label combinations of longevity interventions in the data set of Yen and Mobbs [10].

### Test of superposition principles

To quantify the consistency of the empirical data with the proposed SP’s, we compare the quality of a fit constrained to satisfy a given SP with that of an unconstrained fit. All fits are based on 3-parameter survival functions of the form

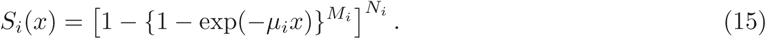

Within reliability theory, the parameters of (15) are interpreted as the failure rate of redundant elements *µ_i_*, the number of redundant elements *M*_*i*_ and the number of essential functional modules *N*_*i*_ [30]. We should like to emphasize, however, that our use of this particular functional form in the present context is motivated solely by the observation that it is sufficiently versatile to provide satisfactory fits to a wide range of empirical survival curves using a moderate number of parameters. The parameters *M_i_* and *N_i_* will therefore not be constrained to take on integer values. To verify that our conclusions do not depend on the particular family of survival functions that is used to implement the analysis, we have carried out a second set of fits using a three-parameter logistic mortality model [24]. The exemplary results shown in the Supplementary Material (fig. S25) are indistinguishable from those based on (15).

The fit algorithm described in the **Methods summary** minimizes the sum of squares of the mean square deviations (SSD) corresponding to the four curves in the quadruple

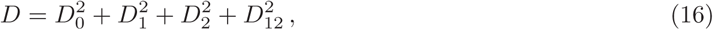

where 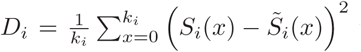 with 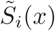 denoting the empirical surviving fraction and *k*_*i*_ the number of data points. In the first step of the analysis the survival curves are fitted individually, which implies that the terms in (16) are independent. In the next step a second fit is carried out under the constraint imposed by the SP of interest. The implementation of this step differs between the different SP’s introduced above.

- For the GM-SP, the survival curves *S*_0_, *S*_1_, and *S*_2_ are represented by three survival functions of the form (15), and the fourth curve *S*_12_ is constructed according to (5) using the numerical computation of inverse functions. The 9 parameters entering the three functions are then adjusted to optimize the fit to the data quadruple.
- By contrast, a direct fitting algorithm constrained to satisfy the CR-SP (3) will in most cases fail to converge to a valid survival curve. This reflects the restrictive conditions on the individual curves imposed by this SP that were exemplified above using the Weibull function (7). To overcome this difficulty, we further constrained the fitting procedure by demanding that the four survival curves in the quadruple take the specific form

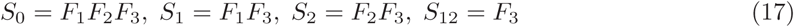

where the *F*_*i*_(*x*) are again represented by three-parameter functions (15). This enforces the validity of the CR-SP (3) but also implies that the curves have to be ordered according to *S*_12_(*x*) ≥ *S*_1_(*x*), *S*_2_(*x*) ≥ *S*_0_(*x*) for all *x*.
- Finally, for the implementation of the GS-SP, the fit determines a single three-parameter survival function *S*_0_(*x*) along with the 4 parameters *b*_1_, *b*_2_, *q*_1_, *q*_2_ entering the scaling transformations (13) and (14).

Note that different quadruples have in general a different inherent difficulty to be fitted. As we are not interested in the absolute quality of the fits provided by the particular function (15) but in the relative quality of the constrained fits associated with different SP’s, we normalize the SSD *D* for fits that fulfill an SP by the SSD *D*_ind_ obtained when the four curves are fitted independently. Doing this enables us to assess how well the different SP’s are satisfied for different quadruples of data. It turns out that all 24 quadruples can be fitted reasonably well by at least one of the three SP’s.

Examples of three experimental quadruples and the corresponding fits are shown in fig. 1. For each column, a different type of SP yields the lowest relative SSD. In column a), the CR-SP provides the best quality of the fit, in column b) it is the GM-SP, and the GS-SP in column c). In all three cases the relative SSD *D*/*D*_ind_ of the best fit is very close to unity, showing that the corresponding SP is satisfied with high accuracy. A full set of figures showing all pairs of empirical survival curves with their respective optimal fits can be found in the Supplementary Material (Figs. S1-S24).

**Figure 1.**
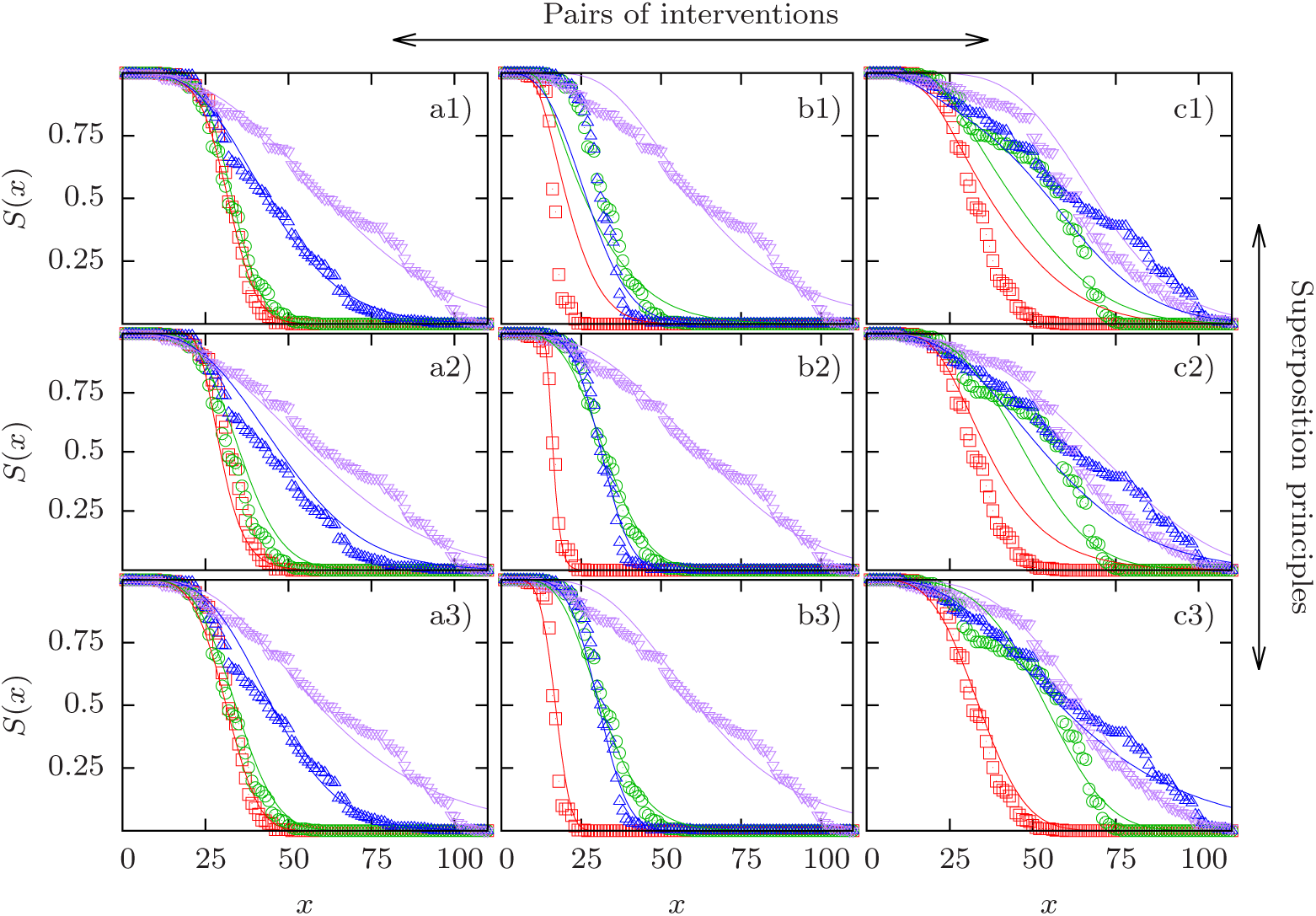
Comparison of experimental survival curves and model fits for three cases. Experimental survival curves are depicted by symbols and their respective fits by lines. Columns correspond to different pairs of interventions and rows correspond to different superposition principles (SP’s). Column a) shows the quadruple 1000–1011, column b) the quadruple 0010–1011, and column c) the quadruple 1010–1111. Row 1) shows the competing risks SP, row 2) shows the generalized multiplicative SP, and row 3) show the generalized scaling SP. Red squares represent the baseline curve, green circles and blue upward triangles display the two single interventions in the order of their position in the binary string (green circles first, blue upward triangles second), and purple downward triangles correspond to the combined interventions. The fits in panels a1), b2) and c3) have the best quality in their respective column in terms of their sum of squared deviations *D* defined in (16). The relative SSD’s of the three best fits are *D*/*D*_ind_ = 0.834 (a1), 1.06 (b2) and 1.06 (c3), respectively.

It is evident that the examples shown in the three columns represent different patterns. Column a) depicts the interaction of the *clk-1* mutation with DR at low temperature. The effect of *clk-1* on life span is hardly detectable in the absence of DR but becomes significant when DR is applied as well, thus providing an example of synergistic epistasis for mean life span between two interventions of widely different individual effects. As we have seen that apparent synergistic epistasis between interventions of strongly unequal effects is a generic feature of the CR-SP, it is not surprising that this SP is able to describe these data very well. By contrast, the survival curves in column b) show two interventions of similar effect (low temperature and DR applied to the *daf-2* mutant) which combine essentially multiplicatively in terms of mean life span. Since the GM-SP satisfies multiplicativity of the median life span by construction, it yields the best fit to the data in this case. Finally, column c) displays a case of apparent antagonistic epistasis, where the combined interventions of the *clk-1* mutation and DR on the background of low temperature and *daf-2* is essentially indistinguishable from the effects of the individual interventions. The GS-SP is the only one of the three SP’s that is principally able to account for antagonistic epistasis for mean life span, and therefore it provides the best description of these data.

The correlation between the preferred SP and the type of epistasis on the level of median life span that is observed in the examples shown in fig. 1 holds quite generally across all 24 pairwise comparisons. In fig. 2 we plot the ratio *D*/*D*_ind_ vs. the epistasis coefficient of median life spans defined as

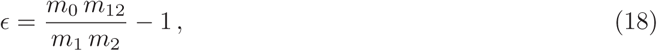

which vanishes under the multiplicative condition (6), and is positive (negative) in the presence of synergistic (antagonistic) epistasis. This figure thus illustrates the relationship between epistasis for median life span quantified by *ϵ*, and epistasis on the level of survival curves quantified by the minimal value of *D*/*D*_ind_. As discussed previously survival curves obeying the CR-SP tend to favor synergistic epistasis for mean life span and hence there is a negative correlation between the epistasis coefficient *ϵ* and the normalized SSD *D*/*D*_ind_ for this SP (red squares in fig. 2). In the same manner, curves obeying the GM-SP display a lower goodness of fit (larger relative SSD) the larger the absolute value 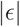. As there is no a-priori preference of the GS-SP for a particular type of epistasis, fits performed under this principle yield decent results for all values of ϵ. Accordingly, looking only at the SP that yields the best result for a given data quadruple, one observes that the CR-SP works best for data with strong synergistic epistasis while GM-SP works best when epistasis is weak. Because both principles perform poorly with strong antagonistic epistasis, the GS-SP yields the lowest SSD in this regime.

**Figure 2.**
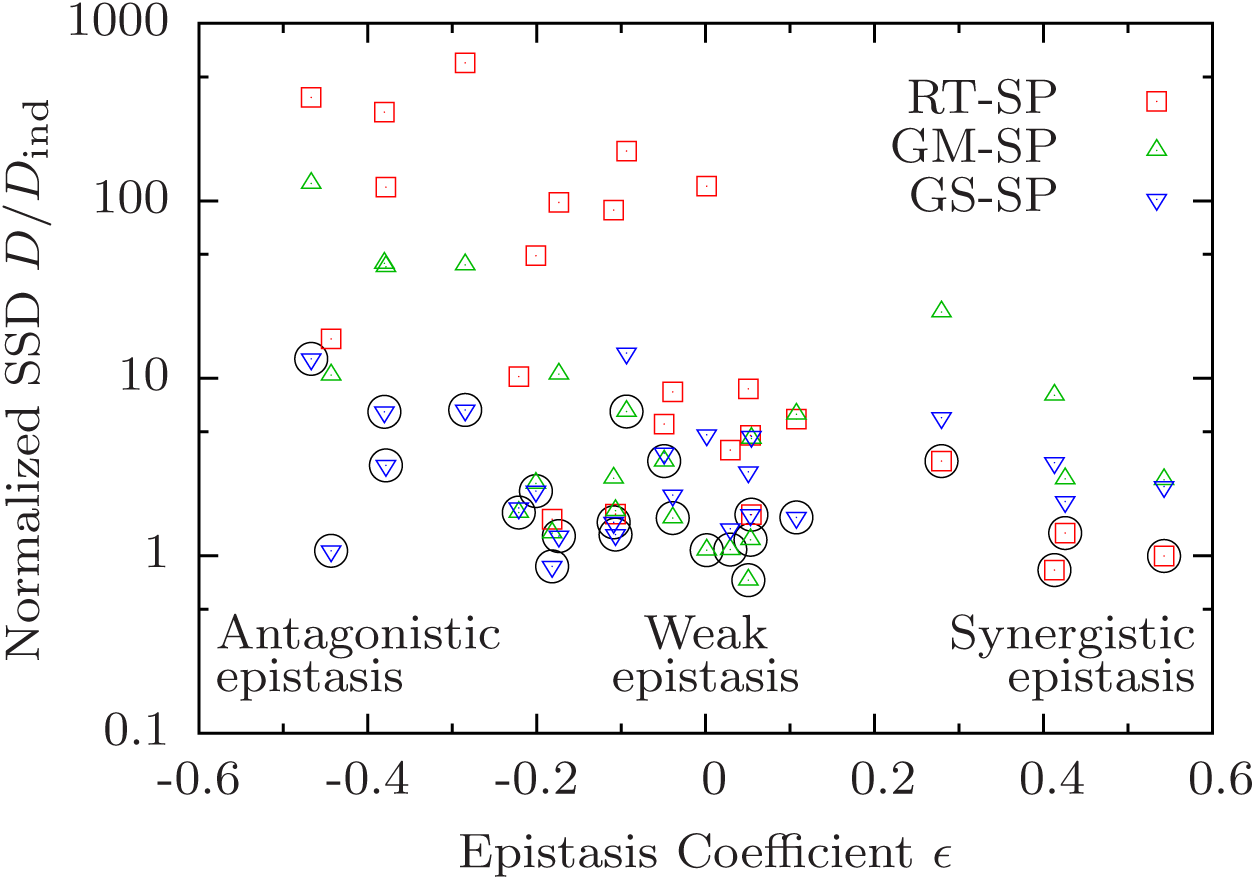
Preference for different superposition principles correlates with epistasis in median life span. The sum of squared deviations *D* of survival curves satisfying a superposition principle (SP) is divided by the SSD *D*_ind_ of independently fitted curves and shown in dependence on the epistasis coefficient *ϵ* defined in (18). Each symbol corresponds to a combination of a data quadruple and a SP. The SP yielding the best result for a given quadruple is marked by a black circle.

Apart from this conspicuous pattern, however, the most striking feature of fig. 2 is that epistasis on the level of survival curves is remarkably weak, in the sense that the minimal value *δ ≡* min_SP_ {*D*/*D*_ind_} is often close to unity. Specifically, *δ <* 2 in 16 out of 24 cases, and there is only one quadruple (0000–1100, see fig. S1) for which *δ* > 10. The latter corresponds to the combination of *daf-2* and low temperature, which has been found in previous work to display significant antagonistic interactions [10]. In fact, three out of the four pairs with *δ >* 5 comprise these two interventions on different backgrounds. On the other hand, the quadruples 1000–1011 (figs. 1a) and S23) and 1010–1111 (figs. 1c) and S20) display significant positive (*ϵ* > 0.4) and negative (*ϵ* < −0.4) epistasis for median life span, respectively, but both have *δ* ≈ 1. Overall, fig. 2 makes it evident that epistasis for median life span is a very poor predictor for the existence of interactions on the level of the survival curves.

## Discussion

The superposition principles introduced above quantify different natural notions of independence between longevity interventions. The GM-SP generalizes the commonly used multiplicative model for relative life span increases to the quantile function *Q*(*s*), which is sufficient to predict the survival curve of the combined intervention from the survival curves representing the individual interventions. The CR-SP follows under rather general conditions from a modular structure of the functions on which the survival of the organism depends, as exemplified by (but not restricted to) the reliability theory of ageing. Finally, the GS-SP is based on the assumption that longevity interventions can be viewed as generalized scaling transformations applied to the survival curve, which are commutative and therefore yield a unique prediction for the combined survival curve.

Two of the three SP’s (GM and CR) are non-parametric, in the sense that they can be formulated without reference to a particular parametrization of the survival curves Si or the longevity transformations *Ti*. One might have expected that this property would facilitate the application of these SP’s to data, but this is in fact not the case. The direct test of the CR-SP is considerably exacerbated by the fact that the insertion of an arbitrary set of survival functions *S*_0_, *S*_1_ and *S*_2_ on the right hand side of (3) does not generally produce a valid survival curve, and similar problems may arise for the GM-SP (2). In comparison, the application of the parametric GS-SP is more straightforward and has the additional benefit of yielding some insight into the nature of the longevity transformations involved through the estimates of the parameters *b*_*i*_ and *q*_*i*_ in (13). As we have outlined above, the CR-SP and the GS-SP have natural interpretations in term of the competing risks, proportionate hazard and accelerated failing rate models of survival analysis [31].

An important conclusion from our approach is that independence of longevity interventions on the level of survival curves does not generally imply the absence of epistasis for mean life span. This point is most clearly illustrated by the CR-SP, which is based on a biologically plausible concept of independence in terms of modularity of vital functions, and implies additivity of age-dependent mortality in the sense of (12). Nevertheless, as we have demonstrated for a class of survival functions, interventions combined according to the CR-SP can display substantial synergistic epistasis in their effect on life span. We believe that this is true irrespective of the specific form of the survival curve, and a proof of this conjecture would be of considerable interest. For the GS-SP we have shown that the apparent epistasis for mean life span can be positive or negative depending on the parameters entering the longevity transformaion.

Our explorative investigation of the empirical data set of [10] shows that all quadruples of survival curves can be fitted rather well by at least one of the SP’s. This indicates that ‘true’ epistatic interactions that would become manifest in a violation of the general superposition principle (4) are rare, even though epistasis for mean life span can be quite significant (see fig. 2). It remains to be seen if this outcome is specific to the data set under investigation. None of the three suggested types of SP’s was found to be universally preferred. Instead, the preference for a given SP is correlated with the amount and sign of epistasis on the level of median life span. In this way our analysis decomposes the 24 pairs of survival curves into three classes with qualitatively different patterns of interactions. So far we have not been able to clearly attribute individual pairs of interventions to specific classes. With one exception (the combination of low temperature and *daf-2*, which always falls into the GS class) the attribution generally varies according to the identity of the two background interventions.

Moreover, despite our pooling of data obtained from different experiments, the attribution appears to be significantly affected by measurement error. This is illustrated by the Supplementary Figure S25, which displays the results of an analysis using a single set survival experiments corresponding to the largest cohort size. Although the overall pattern is similar to fig. 2, the attribution of specific pairs of interventions to their preferred SP’s differs considerably and the correlation with the epistasis parameter *ϵ* is weakened. We expect that the recently developed methods for the generation of high resolution survival curves [33,34] will help to alleviate this problem and allow to extract specific functional information from the kind of analyses proposed here.

## Methods summary

The fitting algorithm aims to minimize the sum of squared deviations *D* defined in (16). Even though the survival curves occurring in this paper have a relatively simple shape, it is still rather difficult to fit several interdependent curves at once. In particular, standard hill-climbing algorithms tend to converge to suboptimal minima of *D*. We therefore use an evolutionary algorithm that consists of the following steps:

1. The algorithm is initialized with a population of *n* quadruples of survival functions. Initial parameter values are *µ*_*i*_ = *M*_*i*_ = *N*_*i*_ = *b*_*i*_ = *q*_*i*_ = 1.0.
2. Next *m* offspring are created that descend from randomly chosen parents. The parameters of the children are equal to the parent’s parameters multiplied with a factor *e*^*uX*^, where *X* is uniform random variable on [−1,1] and *u* > 0 is the mutation strength.
3. Out of the total population of the *n* + *m* individuals, the *n* with lowest SSD survive. These individuals make up the next generation.
4. Mutation strength *u* is decreased by a constant factor, and the algorithm continues with the second step.

For the fits in this paper we chose *n = m* = 180 and ran the algorithm for 2500 generations. We chose *u* = 1 for the initial generation and decreased it in every generation by a constant factor such that *u* = 0.01 in the final generation.

## Acknowledgements

We thank Adam Antebi, Scott Pletcher, Björn Schumacher and Yidong Shen for useful discussions, Kelvin Yen for giving us access to his original data, and an anonymous reviewer for constructive remarks. This work was supported by BMBF within Sybacol. RM acknowledges the DST-INSPIRE faculty scheme, Department of Science and Technology, India for financial support.

## Author contributions

All authors devised the study and carried out mathematical analyses. SN, JN and JK wrote the paper.

